# Event2Flow: Scalable imaging of peripheral and cerebral hemodynamics with event-based vision sensors

**DOI:** 10.64898/2026.02.17.706291

**Authors:** Quanyu Zhou, Weiye Li, Nico Messikommer, Zhenqiang Li, Tian Jin, Xuyang Chang, Baoyuan Zhang, Shiyao Guo, Lin Tang, Michael Reiss, Xiong Dun, Zhenyue Chen, Davide Scaramuzza, Daniel Razansky

## Abstract

Accurate blood flow mapping over mesoscale fields of view is essential for understanding physiological and pathological processes, yet conventional optical methods often rely on bulky high-speed cameras that generate massive datasets with excessive computation burden. Here, we introduce Event2Flow, a compact and data-efficient framework leveraging event-based vision sensors, which asynchronously capture brightness changes with sub-millisecond latency and minimal data redundancy. Event2Flow supports multiple contrast mechanisms for flow measurement, including speckle fluctuation and particle tracking. By correlating the event count with flow velocity through simulations and experiments, we first demonstrate its application in laser speckle imaging for noninvasive mapping of mouse ear vasculature and ethanol-induced hemodynamic changes. When integrated with widefield fluorescence localization microscopy and point spread function engineering, Event2Flow further enables kilohertz-rate particle tracking for rapid 3D velocity quantification in transcranial brain imaging and snapshot flow direction estimations using event polarity. Overall, Event2Flow offers a scalable alternative to conventional high-speed imaging systems for vascular and neuroimaging applications.

## Introduction

Living biological tissues exhibit high metabolic activity, with the vascular network playing a central role in delivering oxygen and nutrients to sustain cellular activity. Accurate reconstruction of the vascular network – both structurally and functionally – is essential for understanding physiological processes across scales, from individual cells to whole organs. However, quantifying blood flow remains inherently challenging due to: (1) the broad dynamic range of flow velocities, spanning from fast arterial and venous flow (>10 mm/s) to slow capillary flow (<1 mm/s)^1,2^; (2) rapid temporal fluctuations; and (3) the vectorial nature of flow, encompassing both magnitude and direction. These complexities pose significant challenges for existing hemodynamic imaging modalities.

Optical imaging is widely utilized in vascular studies due to its high spatial resolution and sensitivity to light-tissue interactions. Most techniques for imaging flow fall into two categories: speckle-based and tracking-based (Fig. 1a). Speckle-based methods, exemplified by laser speckle contrast imaging (LSCI)^3-5^ and optical coherence tomography^6,7^, infer blood flow from temporal fluctuations in speckle patterns caused by red blood cell (RBC) motion. These approaches are label-free and easy to implement but are generally semi-quantitative and lack accurate directional information. Furthermore, conventional intensity-based (frame-based) sensors commonly used in LSCI, such as charge-coupled devices (CCD) and complementary metal-oxide-semiconductor (CMOS) cameras, require fixed exposure time that cannot be altered post-acquisition. The choice of exposure time introduces a tradeoff between sensitivity to slow flow velocities and temporal resolution.

**Figure 1.**
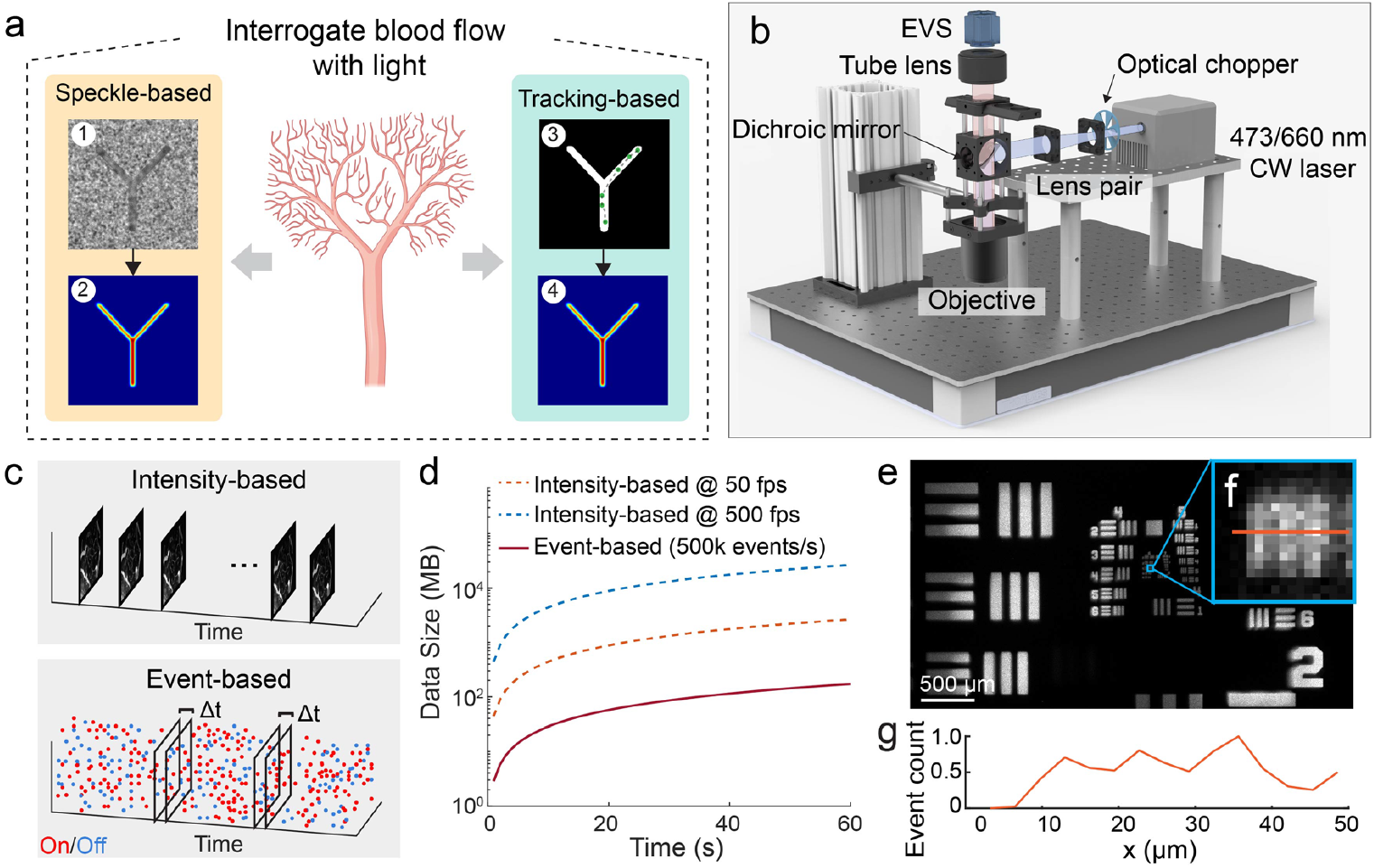
Concept and system design of Event2Flow. **(a)** Overview of classical optical methods for blood flow imaging, primarily categorized into speckle-based and tracking-based approaches. **(b)** Schematic of the Event2Flow system, which integrates an EVS into a customized widefield microscope. **(c, d)** Comparison between an intensity-based camera and an EVS in terms of output format and data size for a fixed acquisition duration. **(e)** FOV and resolution characterization using a USAF resolution target under modulated illumination. **(f)** Zoom-in of Group 6, Element 4 from panel **e. (g)** Line profile along the orange line in **f**. Diagrams in **a** were created using BioRender (https://biorender.com). Optical components in **b** were adapted from 3D models provided by Thorlabs, Inc.

Tracking-based methods offer more direct and quantitative flow readouts. For example, kymography-based methods in laser scanning microscopy track labeled RBCs or plasma markers via line scanning along vessel centerlines^8,9^. RBCs appear as streaks, from which flow velocities could be extracted via computational algorithms^10,11^. However, this approach is inherently restricted to single-vessel measurements, making full-field mapping challenging. To address these limitations, we previously proposed widefield fluorescence localization microscopy (WFLM)^12-14^, which tracks fluorescently labeled particles or cells using classical widefield microscopy. WFLM enables simultaneous structural and functional imaging over a large field of view (FOV), while capturing high-velocity fluorescent targets requires high-speed cameras operating at kilohertz rates. Besides their significant cost and scarce availability, such implementations can generate tens of gigabytes of data per minute that exceed the bandwidth of typical data interfaces (e.g., USB 3.0 or CameraLink), thus necessitating onboard memory buffers and further increasing system complexity, size, and cost, all of which hinder broader adoption.

The need for high-throughput, compact, and cost-effective hemodynamic imaging solutions motivates the exploration of alternative solutions. In typical flow imaging scenarios, vascular targets are spatially sparse, with most pixels in the FOV contribute little information but still consume bandwidth and inflate data volume. Event-based vision sensors (EVSs)^15,16^, also known as neuromorphic cameras, may offer a promising alternative. With sub-millisecond (ms) temporal resolution, EVSs have gained traction in fields such as automotive vision^17-20^, robotics^21,22^, super-resolution microscopy^23,24^, and light field microscopy^25^. Unlike conventional intensity-based cameras, EVSs operate on a bio-inspired event-driven architecture: each pixel responds asynchronously to changes in logarithmic brightness, recording only where intensity increases (“On” event) or decreases (“Off” event).

The discrete, near-binary output of EVSs – containing only timestamp, location, and polarity – may seem incompatible with measuring continuous processes like blood flow. In this work, we bridge this gap by introducing Event2Flow, a high-throughput imaging framework that leverages EVSs for blood flow mapping across both speckle-based and tracking-based contrast mechanisms. We first simulated the EVS response to speckle contrast variations and established a direct correlation between event count and flow velocity. Unlike traditional methods such as LSCI, Event2Flow operates asynchronously at the pixel level and does not require predefined frame rates. Using this approach, we performed real-time noninvasive vascular mapping of peripheral vessels in the mouse ear and assessed flow changes in response to ethanol stimulation. Beyond speckle imaging, we integrated EVSs with WFLM and demonstrated its capability to quantitatively resolve blood velocities at the single-vessel level. Notably, the polarity information in EVS output allows snapshot flow direction estimation without explicit particle tracking, marking a fundamental distinction from previously reported WFLM implementations. Lastly, by incorporating point spread function (PSF) engineering, we extended Event2Flow to three-dimensional (3D) morphological and functional imaging of vascular networks. With significant reduction in system size, cost, and data volume, Event2Flow presents an accessible and scalable solution for vessel-related imaging, facilitating its adoption across diverse applications in vascular research and beyond.

## Results

### Event2Flow system design and imaging performance

The Event2Flow system design is illustrated in Fig. 1b, where an EVS is integrated into a customized widefield microscope. The system incorporates both 660 nm and 473 nm laser sources, enabling flexible switching between LSCI and WFLM imaging modes (see *Methods* for details). Unlike traditional intensity-based cameras that capture full-frame time-lapse images, the EVS records only changes in pixel intensity caused by blood flow (Fig. 1c). Depending on whether the intensity increases or decreases, “On” and “Off” events are recorded, with a temporal latency of ∼100 μs. Each event is encoded as (*x, y, p, t*), where *x* and *y* denote spatial coordinates, *p* indicates event polarity, and *t* is the timestamp. This event-driven architecture leads to 1∼2 orders of magnitude reduction in data volume compared with intensity-based cameras operating at 50 or 500 fps with 8-bit dynamic range and equivalent pixel resolution (Fig. 1d). The FOV and lateral resolution of the system were characterized by imaging a USAF resolution target placed on a fluorescence slide (Fig. 1e-g). Modulated illumination was achieved by placing an optical chopper in the light path, generating fluorescence fluctuations to trigger events. The system provided a FOV of 4.1× 2.3 mm^2^ and was able to resolve Group 6, Element 4 of the USAF target. The corresponding line profile is shown in Fig. 1g, where line pairs with a spacing of 11.05 μm were clearly distinguishable.

### Event2Flow for label-free hemodynamic imaging in the mouse ear via speckle contrast decoding

We first applied Event2Flow for label-free imaging of peripheral vascular in the mouse ear, both under baseline conditions and following ethanol stimulation, utilizing a laser speckle setup with coherent illumination and backscattered light detection (Fig. 2a). The imaging system was configured with 2× magnification to satisfy the Nyquist sampling criterion for speckle imaging, while speckle size was characterized by temporarily replacing the EVS with an intensity-based camera. To understand how EVS responds to flow-induced speckle fluctuations, we simulated both intensity-based and event-based recordings across varying flow velocities using a dynamic laser speckle model^26^ (Fig. 2b). Event2Flow differs from conventional LSCI in two key aspects: (1) LSCI captures blurred speckle patterns due to limited temporal resolution and infers flow from speckle contrast loss, whereas Event2Flow directly detects instantaneous speckle intensity changes at sub-ms temporal resolution; (2) LSCI records full-frame intensity images, while EVS outputs sparse, background-free event streams.

**Figure 2.**
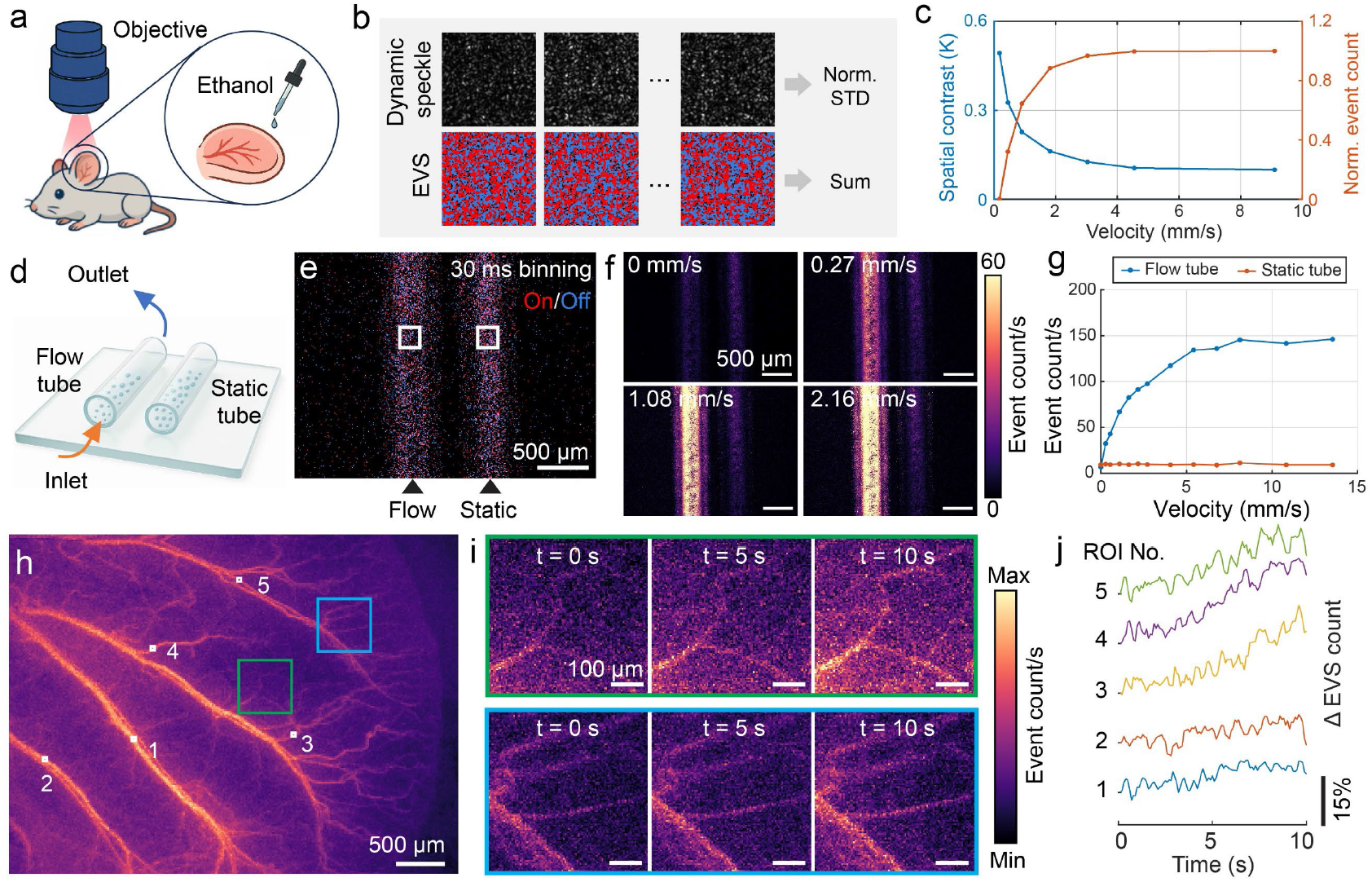
Event2Flow for label-free hemodynamic imaging of the mouse ear. **(a)** Illustration of mouse ear imaging and topical ethanol stimulation. **(b)** Simulated laser speckle evolution and corresponding EVS response. Speckle fluctuations were quantified using normalized standard deviation (for classical LSCI) and summed event count (for Event2Flow), respectively. **(c)** Comparison of spatial speckle contrast and event count as a function of flow velocity. **(d)** Illustration of phantom experiment with two parallel microtubes filled with 1.2% intralipid. The left tube was connected to a syringe pump (flow tube) and the right used as a static reference (static tube). **(e)** Event-rendered image generated by accumulating events over a 30 ms binning window. **(f)** Event count maps acquired at different flow velocities in the flow tube. **(g)** Quantification of event counts at selected regions of interest (ROIs) marked in **e**, plotted as a function of flow velocity. **(h)** Event count map of the mouse ear acquired with 10 s accumulation time. **(i)** Time-lapse event count maps in selected ROIs labeled in **h** following ethanol application, rendered with 0.1 s accumulation time. **(j)** Relative event count changes in selected vessels labeled in **h**, indicating a synchronized increase in blood flow post stimulation.

We hypothesized that faster flow would induce more rapid speckle fluctuations, resulting in a higher event count, which is defined as the number of brightness changes detected within a given time window. Simulations of 5,000 particles moving at preset velocities confirmed a positive correlation between event count and flow velocity (Fig. 2c). As expected, the spatial speckle contrast *K* decreased with increasing velocity, consistent with classical LSCI principle. To experimentally validate this, we conducted phantom tests using two parallel microtubes filled with 1.2% intralipid to generate dynamic speckle contrast (Fig. 2d). One tube was connected with a syringe pump to produce variable flow inside (“flow tube”), while the other serves as a static reference (“static tube”). An event-rendered image generated by accumulating events over a 30 ms binning window is shown in Fig. 2e. Event count maps showed increased counts in the flow tube as velocity increased, while the static tube exhibited a relatively constant value (Fig. 2f, g). The existence of event count in the static tube is likely due to Brownian motion of intralipid molecules. At higher velocities (> 5 mm/s), the event count exhibits a saturating trend, likely due to the speckle correlation time becoming much shorter than the EVS latency (Fig. 2g).

We then imaged the mouse ear *in vivo*, where reported blood flow velocities typically range from 0 to 6 mm/s^27,28^. A 10 s accumulation of events yielded a high-contrast vascular map, with event counts decreasing from large vessels to smaller branches (Fig. 2h), confirming that event count is sensitive to blood flow. Beyond static imaging, high temporal resolution is crucial for capturing dynamic changes in response to external stimuli. We therefore investigated how accumulation time impacts the image quality of the vascular map. The contrast-to-noise ratio (CNR) increased rapidly with short accumulation time and then plateaued at longer accumulation time. Based on that, we selected 0.1 s accumulation time for stimulation experiments, balancing temporal resolution and image quality (CNR = 3.36). Following topical application of ethanol to the mouse ear, we observed a rapid increase in event count in selected regions (Fig. 2i), consistent with the known vasodilatory effects of ethanol^29^. Temporal profiles of relative event count changes revealed a global increase in blood flow (Fig. 2j), aligning well with results obtained from a commercial LSCI system, but with a tenfold improvement in temporal resolution. These findings validate event count as a label-free and computationally efficient metric for *in vivo* hemodynamic mapping.

### Event2Flow for transcranial single-vessel-level flow mapping in the murine brain via 2D particle tracking

While Event2Flow has proven effective for peripheral vessel imaging by leveraging speckle fluctuations, quantifying flow in larger vessels (flow speed > 5 mm/s) and deeper tissues remains challenging due to complex light scattering and overlapping vascular structures. To achieve higher spatial sparsity and enable single-vessel-level flow quantification, we applied Event2Flow to track fluorescent particles in the murine brain (Fig. 3a). Unlike traditional intensity-based cameras that yield images with high dynamic ranges, EVSs generate sparse “On” and “Off” events triggered by local brightness changes induced by particle motion (Fig. 3b).

**Figure 3.**
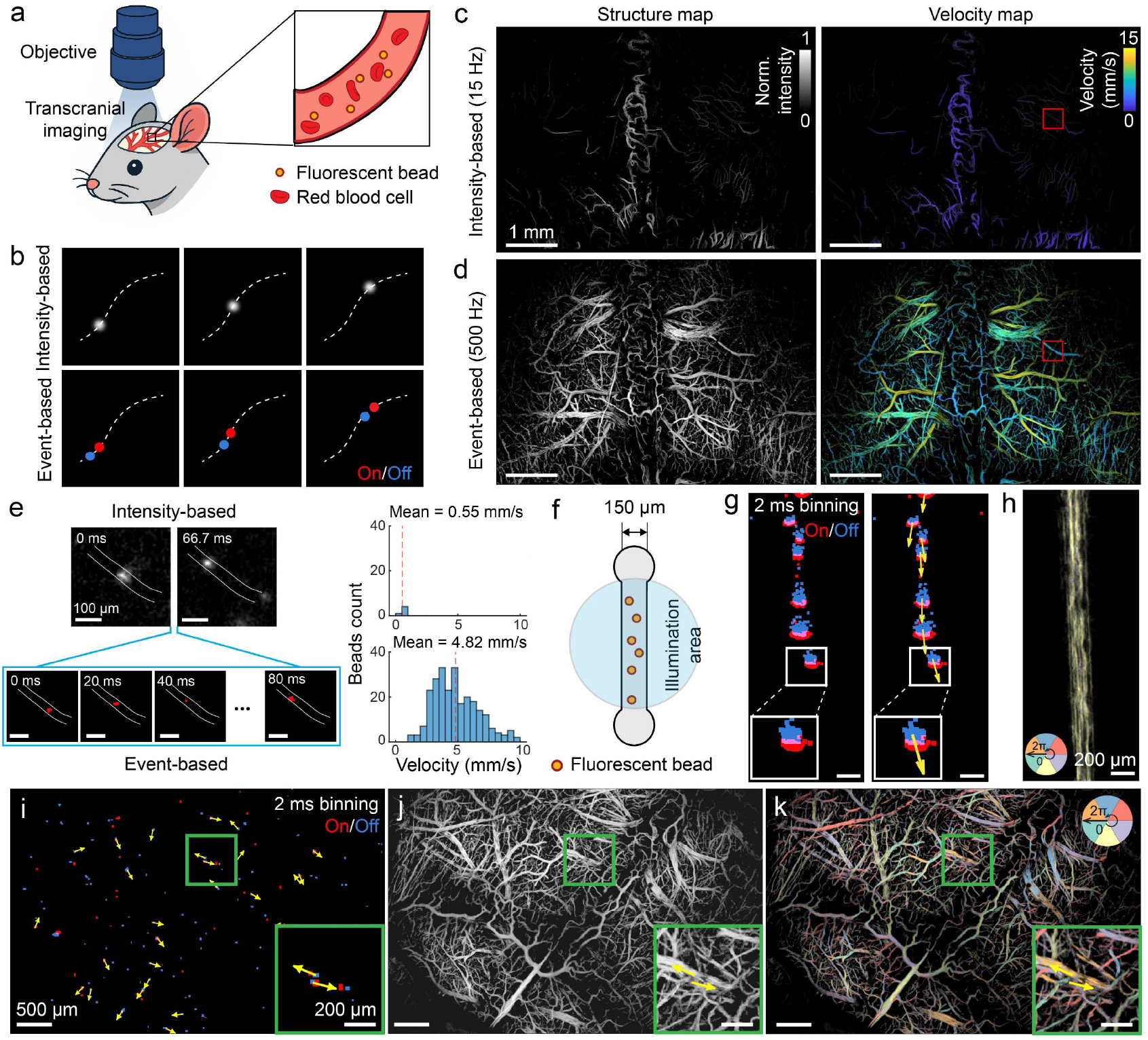
Event2Flow for transcranial structural and functional imaging of murine brain. **(a)** Illustration of transcranial murine brain imaging based on fluorescent particle tracking. **(b)** Comparison between intensity-based and event-based recordings. **(c, d)** Reconstructed structural and flow velocity maps from the intensity channel and event channel recording of a hybrid vision sensor. **(e)** Time-lapse images of a selected skull vessel (labeled with red squares in **c** and **d**), showing reduced motion blur in the event channel. Right: corresponding velocity histograms. **(f)** Illustration of a phantom experiment for snapshot flow direction estimation using event polarity, involving a straight microtube with flowing particles. **(g)** Event-rendered image with a 2 ms binning window revealed spatially paired “On” and “Off” events. Flow direction vectors (yellow arrows) are defined from “Off” to “On” events. **(h)** Flow direction map reconstructed by aggregating flow vectors over continuous recording. **(i-k)** Event-rendered image with a 2 ms binning window from transcranial murine brain recordings, with tracking-free reconstruction of structural (**j**) and flow direction (**k**) maps. Zoom-in views (green squares) highlight opposing flow directions in neighboring artery and vein.

We used a hybrid vision sensor to simultaneously record intensity frames at 15 fps (“intensity channel”) and event streams at 500 fps (“event channel”), allowing direct comparison between conventional and event-based readouts. Following intravenous injection of fluorescent particles, we recorded for 5 mins and performed particle localization and tracking using an open-source TrackNTrace toolbox^30^ (see *Methods* for details). Structural and flow velocity maps were reconstructed from both the intensity channel (Fig. 3c) and event channel (Fig. 3d). For event channel reconstruction, only “On” events were used due to their shorter latency relative to “Off” events. The intensity channel primarily captured slow-moving particles in skull vessels, whereas the event channel revealed both skull and brain vessels across a broader velocity range, leveraging its low-latency detection capability. This contrast is further confirmed with time-lapse images and velocity histograms from a representative skull vessel (Fig. 3e), as labeled in Fig. 3c, d. Due to the limited frame rate of the intensity channel, only low-speed particles could be reliably tracked (mean velocity = 0.55 mm/s), while fast-moving particles exhibited large inter-frame displacements, resulting in motion blur and tracking failure. In contrast, the event channel resolved a higher mean velocity of 4.82 mm/s. We also noticed missing low-velocity readouts in the event channel, likely because particles moving at slower speeds (e.g., 1 mm/s) produce sub-pixel displacements of ∼2 µm at 500 fps acquisition, which are insufficient to trigger events. These findings underscore the complementary strengths of hybrid sensors in capturing a broader dynamic range of flow velocities.

In addition to the low latency and data efficiency, a unique advantage of EVS lies in its ability to record both event polarities: “On” events are triggered when a particle arrives at a location at the current time point, while “Off” events correspond to their previous positions. This dual-polarity information enables flow direction estimation without the need for tracking – simply by identifying the spatial relationship between paired “Off” and “On” events. The flow direction is inferred as a vector pointing from “Off” to “On” event. We first validated the idea in a straight microtube with fluorescent particles flowing inside (Fig. 3f). The event-rendered image with a 2 ms binning window revealed spatially paired “On” and “Off” events (Fig. 3g, left). By identifying and linking these pairs, we estimated local flow vectors (Fig. 3g, right), and subsequently generated a complete flow direction map by aggregating vectors across a 30 s recording (Fig. 3h). We then applied this method for transcranial recordings in the murine brain. The event-rendered image with a 2 ms binning window is shown in Fig. 3i. Integration over a 5 min recording yielded a super-resolved vascular map and color-encoded flow direction map, reconstructed without tracking (Fig. 3j, k). The resulting direction map exhibited symmetrical flow patterns across hemispheres and inverse flow directions in paired arteries and veins, as illustrated in the zoom-in views (Fig. 3i-k), validating the accuracy of polarity-based flow direction estimation.

### Event2Flow for depth-resolved microcirculation flow mapping in murine brain via PSF engineering

To extend Event2Flow beyond planar (2D) imaging and enable depth-resolved measurements, we integrated PSF engineering into the system by encoding depth information in the PSF geometry. The absence of intensity information in EVSs poses a challenge for most PSF engineering strategies, such as astigmatism^14,31^ and Tetrapod PSF^32,33^, which rely on intensity-based fitting to a numerical PSF model. To address this, we integrated a double-helix (DH) phase mask into the Event2Flow system, which encodes depth as the rotation angle of a dual-lobe PSF, independent of absolute PSF size. The combination of PSF engineering with EVS eliminates the need for axial scanning in conventional fluorescence microscopy, thereby further compressing the data size by encoding the 3D information directly into event stream. In addition, the high acquisition rate of EVS minimizes motion blur from moving particles, thereby reducing its influence on angle estimation. The only hardware modification required was the insertion of the DH phase mask between the objective and tube lens (Fig. 4a). The mask was designed based on an iterative optimization framework^34^ and optimized for a central wavelength of 700 nm. Its broad spectral bandwidth also accommodates the fluorescence emission spectrum of the beads used.

**Figure 4.**
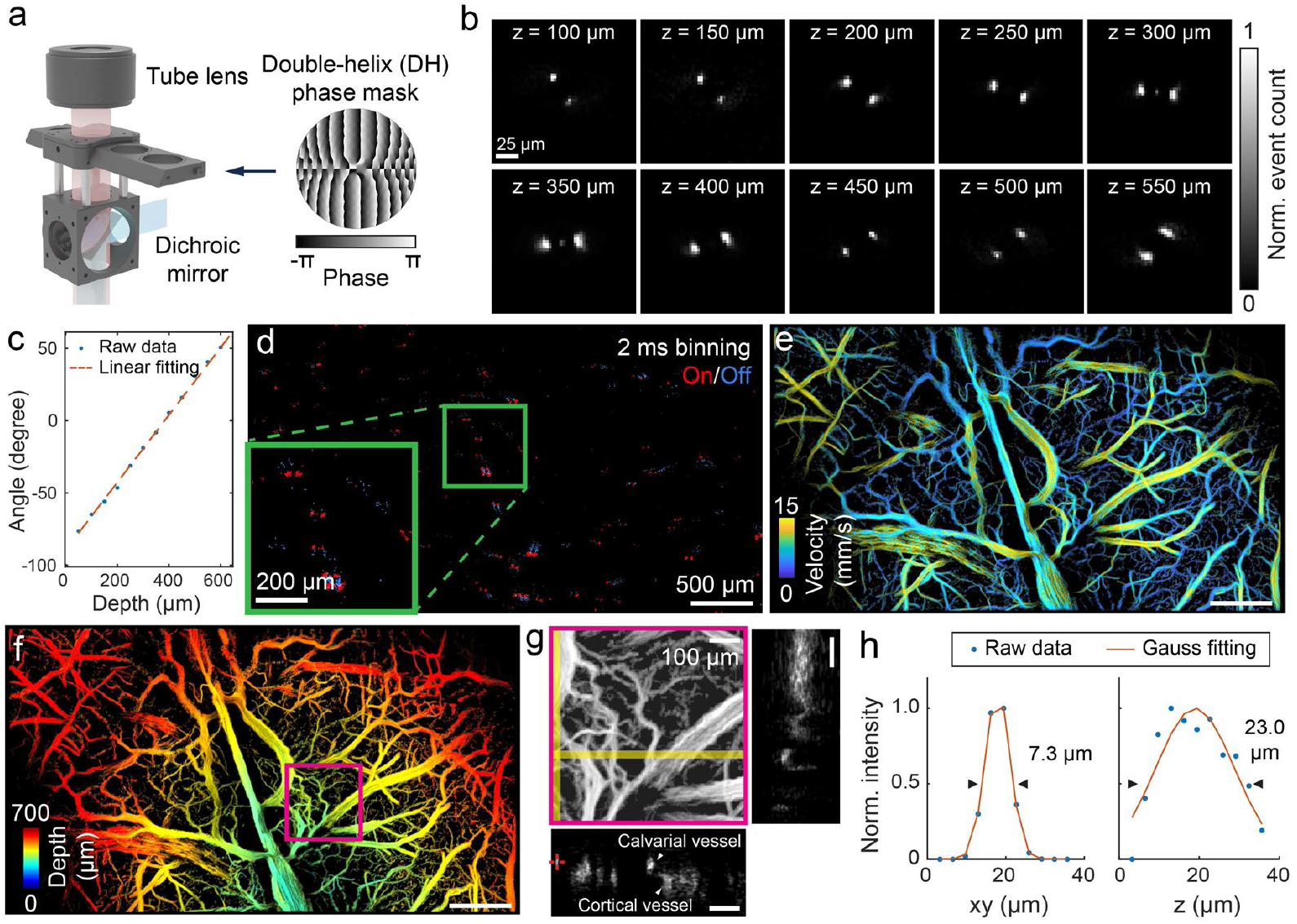
Event2Flow for 3D transcranial microcirculation mapping of murine brain. **(a)** Illustration of 3D Event2Flow system enabled by the insertion of a custom-design DH phase mask. **(b)** Event-rendered PSFs of a single fluorescent bead at varying z positions, showing the dual-lobe feature of DH PSF. **(c)** Measured rotation angle of DH PSF as a function of depth, demonstrating a linear angle-depth relationship. **(d)** Event-rendered image of the murine brain with a 2 ms binning window following beads injection. Inset: zoom-in view of the region marked with the square. **(e, f)** Color-encoded flow velocity and depth maps reconstructed from 3D particle trajectories. **(g)** Zoom-in view of ROI marked in **f**, with side-view projections and selected slices highlighted. **(h)** Lateral and axial line profiles of the vessel labeled in **g**, fitted with Gaussian functions to determine FWHM values.

To generate an angle-depth calibration curve (look-up table), we imaged a single fluorescent bead while incrementally shifting the sample along the z axis under modulated illumination to trigger events. Event-rendered PSFs exhibited the expected dual-lobe rotation with increasing depth (Fig. 4b), and the measured rotation angles displayed a linear relationship with axial displacement across a 600 µm range (Fig. 4c), yielding an average angular sensitivity of 0.23°/μm with R^2^ = 99.78%. We next imaged the murine brain transcranially following intravenous injection of fluorescence beads. A representative event-rendered image with a 2 ms binning window revealed paired spots corresponding to the DH PSF (Fig. 4d). Using their rotation angles and the look-up table, we computed the 3D coordinates of individual particles, followed by particle tracking. Color-encoded depth and velocity maps were then generated by averaging these values across pixels (Fig. 4e, f). Owing to the natural curvature of the skull, the depth map revealed a gradient from the center to the periphery. A magnified region of depth map is shown in Fig. 4g, with xz and yz projections from selected slices revealing two distinct vascular layers – calvarial and cortical vessels – separated by ∼100 μm. A line profile across a representative vessel yielded a full-width-at-half-maximum (FWHM) of 7.3 μm laterally and 23.0 μm axially following Gaussian fitting, reflecting the practical spatial resolution. This depth-resolved capability is particularly valuable in neurovascular studies that require layer-specific analysis of vascular structure and function.

## Discussion

In this work, we introduce Event2Flow, an optical framework that leverages EVSs for blood flow mapping across multiple contrast mechanisms. Replacing traditional intensity-based sensors with EVS offers several key advantages. For speckle-based imaging, Event2Flow enables real-time acquisition and introduces event count as a direct, computationally efficient metric for label-free hemodynamic imaging. For particle tracking-based imaging, Event2Flow eliminates the need for high-speed intensity-based camera with onboard memory. Instead, we utilized sparse events generated by flowing fluorescent beads to achieve single-vessel-level velocity/direction measurements transcranially in the murine brain, which significantly reduced system cost, size, and data load. Furthermore, the inherent polarity information in EVSs (“On” and “Off” events) uniquely enables snapshot estimation of flow direction without relying on particle tracking algorithms. To extend Event2Flow into 3D, we demonstrated its compatibility with DH PSF engineering, requiring only minimal hardware modifications. The DH PSF design encodes depth into the rotation angle of the PSF, making it well-suited for the near-binarized outputs of EVS. This enables 3D visualization of the cerebrovascular network, offering both structural and functional insights.

In addition to optical approaches, hybrid opto-acoustic (OA) modalities and ultrasound imaging have also been employed for blood flow measurements. OA methods can estimate flow velocity by tracking the motion of individual RBCs or cell clusters through decorrelation analysis of repeated A-line acquisitions^35-37^, detecting the displacement of inhomogeneous RBC distribution^38^, or via localization of sparsely-distributed extremely absorbing microdroplets^39^. Additionally, ultrasound localization microscopy^40-43^ offers full-field, single-vessel-level measurements of flow velocity and direction in deep tissues by tracking injected microbubbles at kilohertz rates. These methods typically offer inferior spatial resolution to microscopic optical approaches and further require contact-based detection for ultrasound coupling as well as complex optoelectronic hardware, which can be costly and technically demanding. In contrast, Event2Flow provides an accessible and cost-effective solution, capable of converting any standard widefield microscope into a blood flow imaging platform.

Nevertheless, Event2Flow also comes with its own limitations. First, using event count to infer flow velocity is influenced by factors such as the scatterer density, similar to classical LSCI methods^44^. As a result, event count is better suited for assessing relative flow changes within individual vessels rather than providing absolute flow measurements. Quantitative blood flow measurement would require detailed modeling of speckle decorrelation and camera response, accounting for additional factors from both the sample and camera (i.e., response threshold, noise level). Second, the lack of intensity information restricts sub-pixel localization accuracy of flowing particles, which is commonly achieved vis Gaussian fitting with intensity-based sensors. This limitation affects both the spatial resolution and the ability to quantify slow velocities (i.e., < 1 mm/s), which depends on the minimal detectable displacement between consecutive frames. These challenges could be alleviated by incorporating a hybrid vision sensor, as showcased in this work. Alternatively, subsampling strategy could amplify inter-frame displacements of low-velocity beads through increased accumulation time. Another constraint is the reduced event sensitivity under low-light conditions. Lowering the event detection threshold helps to capture more events but also raises the noise level. This tradeoff limits the utility of EVS for imaging weakly fluorescent targets, such as stained cells, and underscores the need for hardware improvements to enhance event detection sensitivity.

Looking forward, Event2Flow could be adapted for miniature head-mounted microscopes (miniscopes), particularly for imaging in awake, behaving animals – scenarios where data bandwidth and the requirement of system compactness make high-speed cameras impractical. EVSs are expected to overcome this bottleneck by enabling higher acquisition rates without excessive data load. Moreover, the current post-processing pipeline based on open-source localization and tracking software, could be streamlined using deep learning models that convert event data directly into real-time velocity and direction maps. This would improve experimental feedback and facilitate rapid targeting of regions of interest for researchers.

In summary, Event2Flow represents a compact, cost-effective, and data-efficient solution for flow imaging, that supports a variety of contrast mechanisms and imaging scenarios. Its flexible architecture and minimal hardware requirements make it a promising tool for widespread adoption in vascular and neuroimaging communities.

## Methods

### Event2Flow setup

The Event2Flow setup (Fig. 1b) adopts a classical epi-illumination microscopy configuration. The illumination light path integrated two continuous-wave (CW) lasers: a 660 nm laser (gem 660, Laser Quantum, USA) for speckle imaging, and a 473 nm laser (FPYL-473-1000-LED, Frankfurt Laser Company, Germany) for localization experiments. The beam size and converge angle were adjusted with a lens pair (ACN254-050 and AC254-200, Thorlabs, USA).

For speckle imaging, reflected light was separated using a 50/50 beamsplitter (BSW29R, Thorlabs, USA) and collected with a macro lens (Laowa Venus 60 mm, Laowa, China), and focused onto an EVS (EVK4, 1280×720 pixels, 4.86 μm pixel size; Prophesee, France). The system was configured with 2× magnification to fulfill the Nyquist sampling criterion for speckle imaging.

For localization experiments, the illumination beam passed through a scan lens (CLS-SL, EFL=70 mm, Thorlabs, USA), serving as the objective to achieve a wide FOV. Backscattered fluorescence was collected by the same objective, passed through a dichroic mirror (FF495-Di03-25×36, Semrock, USA), and then focused onto the detector via a camera lens (AF Micro-Nikkor 105 mm lens, Nikon, Japan). Data acquisition was conducted using both a hybrid vision sensor (ALPIX-Eiger, 4896×3672 pixels, 1.89 μm pixel size; Alpsentek, Switzerland) and a pure EVS (EVK4, Prophesee, France). The hybrid sensor captured intensity frames at 15 fps (exposure time set to 10 ms) and event-rendered frames at 500 fps, while the pure EVS output raw event streams in the format (x, y, p, t).

### System characterization

The lateral resolution and FOV of Event2Flow system, configured for localization experiments, were characterized by imaging a negative USAF resolution target (R1DS1N, Thorlabs, USA) placed on a fluorescence slide (FSK, Thorlabs, USA) under 473 nm illumination. An optical chopper (MC1F10, Thorlabs, USA) was positioned at the laser output to introduce temporal intensity fluctuations and trigger events.

To measure the speckle size, the EVS was temporarily replaced with a intensity-based camera (a2A2590-60umBAS, 2592×1944 pixels, 2 μm pixel size, Basler AG, Germany). Acquired speckle images were analyzed using spatial autocorrelation. The speckle size on the sensor was defined as the FWHM of the central peak in the autocorrelation map, determined by Gaussian fitting of the central line profile.

### Dynamic laser speckle simulation

Time-lapse laser speckle patterns were simulated using an open-source dynamic speckle evolution model^26^, modified to replace Brownian motion with constant-speed motion in randomized directions. The particle velocity *v* was determined by the speckle correlation time *τ*_*c*_, using the relation:

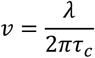

where *λ* is the wavelength. A total of 5,000 particles were initialized at random positions and updated based on the pre-defined velocity. Speckle size and speckle-to-pixel ratio were configured to match the experimental system. The temporal evolution of the speckle pattern was simulated with a time step of 100 μs over 100 steps. Spatial contrast *K* was quantified as the standard deviation divided by the mean intensity of the frame accumulated over the simulation period. To simulate the response of EVS, intensity differences between adjacent frames *I*_*i*+1_ and *I*_*i*_ were computed, where *i* is the frame index. An event was triggered whenever the logarithmic intensity change exceeded a defined threshold *C*, set to 0.22, according to the condition:

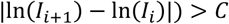

The total event count was obtained by summing all “On” events across the entire FOV over the simulation period.

### Laser speckle contrast imaging

A commercial LSCI system (FLPI, Moor Instrument, UK) was used to record ethanol-induced hemodynamic changes in the mouse ear, with the frame rate of computed perfusion image set to 1 s. Image processing was performed using MoorFLPI software (moorFLPI Review V6.0, Moor Instrument, UK) and custom scripts with MATLAB (MathWorks, MATLAB R2024a, USA) for data visualization and analysis.

### Animal

#### Animal model

All animal experiments were performed in accordance with the Swiss Federal Act on Animal Protection and approved by the Cantonal Veterinary Office Zurich. A total of N = 3 Crl:NU(NCr)-Foxn1^nu^ mice and N = 1 C57BL/6J mouse (10 weeks old, female, Charles River Laboratories, Germany) were used for ear imaging following ethanol stimulation with Event2Flow and LSCI, respectively. For 2D and 3D localization experiments, N = 3 nude mice and N = 2 C57BL/6J mice (9-23 weeks old, female, Charles River Laboratories Inc., USA) were used. The mice were housed in ventilated cages with ad libitum access to food and water. The housing room was maintained was maintained at 22 °C temperature, ∼50% relative humidity, and a 12-hour light/dark cycle.

#### Anesthesia and surgery

Mice were anesthetized with inhalation isoflurane (4% for induction, 1.5% for maintenance) delivered in a gas mixture of oxygen (0.2 L/min) and medical air (0.8 L/min). During surgery and imaging, mice were positioned on a feedback-regulated heating system (PhysioSuite, Kent Scientific, USA) to maintain body temperature at 37 °C, with head-fixed with a stereotaxic frame (Narishige International Limited, UK). Ear imaging was performed in a non-invasive manner. For brain imaging, the mouse scalp was carefully removed 30 minutes after subcutaneous administration of analgesics (Buprenorphine, 0.1 mg/kg). The exposed skull was then moistened using ultrasound gel (Aquasonic Clear, Parker Laboratories Inc., USA). A catheter filled with phosphate buffered saline (PBS) was inserted into the tail vein for injection prior to imaging.

#### In vivo imaging

Ear imaging was performed without external contrast agents. Event data were streamed following topical application of 100 μL 100% ethanol to the mouse ear. For brain imaging, event data were recorded following intravenous injection of 100∼150 μL orange-yellow fluorescent microspheres (1∼5 μm FMOY, Cospheric, USA).

### Data analysis and statistics

#### Speckle analysis

Raw event flow was directly accumulated over fixed time windows to form event count maps. A 3×3 median filter was applied to smooth the images.

#### 2D/3D localization-based image reconstruction

Only “On” events were used for localization-based imaging reconstruction. For data with the hybrid sensor, event-rendered images were spatially binned 2×2 to improve the spatial continuity of individual events. A background activity (BA) filter^45^ was then applied, using a 4×4 spatial window and a 5-frame temporal window, with an empirically determined threshold of 3 to eliminate random, noise-like events. For data acquired with pure EVS, the raw event streams were accumulated with a 2 ms binning window to form a time-lapse image sequence. Particle localization and tracking were performed using the open-source TrackNTrace toolbox^30^, originally designed for super-resolution microscopy. The toolbox outputs trajectories of detected particles, including their spatial coordinates and frame indices. Final localization images were rendered by superimposing all trajectories, while flow velocity maps were computed by averaging the velocities of particles traversing each pixel. For 3D localization, detected spots were grouped as DH PSF if their spatial separation was less than 20 pixels. Depth information was extracted by calculating the PSF rotation angle and referencing a pre-established look-up table. With both the 3D coordinates of each particle and their tracking trajectories, 3D localization and velocity maps were rendered using procedures similar to those described above.

#### Snapshot flow direction map reconstruction

Both “On” and “Off” events were used to estimate the instantaneous flow direction of individual particles. Particles were localized separately in images rendered from “On” and “Off” events. Detected spots were grouped if they were within 10 pixels from each other. The flow direction was defined as the vector pointing from the “Off” event to the paired “On” event. A tracking-free flow direction map was generated by averaging the local flow directions across particles flowing through each pixel location.

## Data availability

The main data supporting the findings of this study are available within the main text. The raw datasets are available for research purposes from the corresponding author upon request.

## Code availability

Localization and tracking of fluorescence emitters were performed with the open-source TrackNTrace toolbox^30^.

## Acknowledgements

The authors acknowledge support from the Swiss National Science Foundation (grants 310030_192757 and 10.003.762).

## Author contributions

Q.Z. conceived the experimental design. Q.Z., W.L., N.M., Z.L., T.J., and L.T. carried out the experiments. Q.Z. and X.C. conducted data analysis. M.R. assisted with the animal experiments. B.Z., S.G., and X.D. contributed to the design of the phase mask. Z.C., X.D., and D.S. contributed to the interpretation of the results. D.R. secured funding and supervised the work. Q.Z. drafted the manuscript. All authors reviewed the manuscript.

